# Genomic epidemiology of *Neisseria gonorrhoeae* isolates from the G-ToG clinical trial, 2014-2016

**DOI:** 10.64898/2026.03.20.713202

**Authors:** Samantha A McKeand, Michelle J Cole, Jeffrey A Cole, Amanda E Rossiter-Pearson, Jonathan D C Ross, Christoph M Tang

## Abstract

*Neisseria gonorrhoeae* remains a major public health threat due to rising incidence and increasing antimicrobial resistance (AMR). To characterise the landscape of gonococci circulating in England a decade ago, we analysed whole genome sequences and clinical metadata from 354 isolates collected during the G-ToG clinical trial between 2014 and 2016. Comparison with global datasets demonstrated that the G-ToG isolates captured the breadth of UK gonococcal diversity, mapping across multiple lineages without evidence of marked bias related to participant demographic characteristics. The phylogenetic structure of gonococcal isolates within the collection was strongly shaped by sexual network, with statistically significant clustering of isolates circulating within heterosexual individuals compared with the men who have sex with men (MSM) transmission group. MSM-associated lineages exhibited higher minimum inhibitory concentration (MICs) to azithromycin and ceftriaxone and were enriched for corresponding chromosomal AMR determinants and AMR-associated plasmids. These data provide a valuable baseline preceding the implementation of several major UK public health interventions, including HIV pre-exposure prophylaxis (HIV PrEP), doxycycline post-exposure prophylaxis (Doxy-PEP) and vaccination with Bexsero. Continued integration of genomic and epidemiological surveillance will be essential to monitor whether these interventions reshape gonococcal transmission dynamics and AMR trajectories, guiding future control strategies.

## INTRODUCTION

*Neisseria gonorrhoeae*, the causative agent of gonorrhoea, remains a major global public health concern, with an estimated 82 million new infections annually (WHO, 2025). Infections commonly present as urethritis in men but are frequently asymptomatic in women, and can lead to severe reproductive complications, including pelvic inflammatory disease, ectopic pregnancy and infertility, if left untreated (Whelan *et al*., 2021). Infection with *N. gonorrhoeae* also increases HIV acquisition and transmission (Galvin and Cohen, 2004).

Increasing antimicrobial resistance (AMR) threatens the effectiveness of the current first-line treatment, ceftriaxone, as well as second-line agents that include azithromycin, ciprofloxacin and cefixime. Most resistance arises through chromosomal mutations (Sanchez-Buso *et al*., 2021); the *penA* (NEIS1753) gene encodes penicillin-binding protein (PBP) 2 and mosaic *penA* alleles, generated through recombination with commensal *Neisseria* spp., reduce binding affinity of β-lactams, particularly extended-spectrum cephalosporins (ESCs). The *ponA* (NEIS0414) gene encodes PBP1, with an L421P substitution reducing penicillin susceptibility and augmenting the effects of altered PBP2. Mutations in the *mtrR* (NEIS1635) promoter (−35A deletion) or coding sequence (A39T and G45D) de-repress expression of the MtrCDE efflux pump, increasing minimum inhibitory concentrations (MICs) to macrolides, penicillin, and tetracycline. Additionally, *porB* (NEIS2020) substitutions at positions G120 and A121 alter porin permeability, reducing influx of hydrophobic antibiotics. Finally, the mutations A2047G or C2599T in 23S ribosomal RNA (rRNA) reduce binding of macrolides, such as azithromycin, with resistance level depending on the number of mutated rRNA copies.

In addition to chromosomal mechanisms, *N. gonorrhoeae* can acquire AMR through plasmids (Cehovin *et al*., 2020). Plasmids are important vehicles for horizontal gene transfer in bacteria, and frequently harbour genes encoding virulence factors, AMR genes and properties that allow bacteria to survive in diverse niches (Slater *et al*., 2008). *N. gonorrhoeae* can harbour three plasmids. The β-lactamase plasmid (p*bla*), which encodes a TEM-type β-lactamase conferring high-level penicillin resistance (Elsener *et al*., 2025) and a conjugative plasmid (pConj), that frequently carries the *tetM* gene, which encodes a ribosomal protection protein that confers high-level tetracycline and doxycycline resistance (Cehovin *et al*., 2020). Finally a cryptic plasmid (pCryp) present in ∼95% of isolates with no known function (Roberts, 1989).

To help combat the issue of AMR and inform national treatment guidelines, the 2014-2016 G-ToG trial assessed the effectiveness of gentamicin as an alternative to the first-line treatment at the time, ceftriaxone (both combined with azithromycin), for the treatment of gonorrhoea (Ross *et al*., 2019). A total of 354 *N. gonorrhoeae* isolates were collected from 296 participants, alongside detailed behavioural and demographic metadata. Combining this metadata with whole genome sequencing (WGS) of isolates provides a valuable opportunity to examine the population structure of *N. gonorrhoeae* circulating in England at that time, characterise transmission networks and investigate the distribution of AMR determinants at high resolution.

## MATERIALS AND METHODS

### Isolate collection

Between October 2014 and November 2016, 720 participants were recruited to the G-ToG clinical trial (Ross *et al*., 2019), across 14 sexual health clinics in England. Adults aged 16-70 were eligible for participation if they had a diagnosis of uncomplicated genital, rectal or pharyngeal gonorrhoea infection, defined by a positive nucleic acid amplification test (NAAT). During the trial, 367 *N. gonorrhoeae* isolates were collected from 307 participants from various anatomical sites, as well as metadata including gender, age, ethnicity and geographical location (Table 1). The trial was registered under International Standard Randomised Controlled Trial Number ISRCTN51783227, and ethical approval was obtained from the Health Research Authority South Central–Oxford C Research Ethics Committee (14/SC/1030).

**Table 1.** Demographic characteristics of participants and associated *N. gonorrhoeae* isolates obtained during the G-ToG trial.

|  |  |
| --- | --- |
| **No. of sequenced isolates** | 354 |
| **No. of participants** | 296 |
| **No. of participants with multiple isolates** | 49 (16.5%) |
| Two isolates | 40 (13.5%) |
| Three isolates | 9 (3.0%) |
| **Gender (male)** | 241 (81.4%) |
| **Median age (range)** | 27 (16-69) years |
| **Clinic location** |  |
| Birmingham | 89 (30.1%) |
| Brighton | 4 (1.4%) |
| Leeds | 39 (13.2%) |
| London | 94 (31.8%) |
| Manchester | 9 (3.0%) |
| Reading | 9 (3.0%) |
| Sheffield | 43 (14.5%) |
| Southampton | 5 (1.7%) |
| **Anatomical sample site** |  |
| Urethra | 165 (46.6%) |
| Rectum | 84 (23.7%) |
| Pharynx | 52 (14.7%) |
| Cervix | 42 (11.8%) |
| Unknown | 11 (3.1%) |

Antimicrobial sensitivity of isolates was determined at the UK Health Security Agency (formerly Public Health England) (Cole, 2020) with MICs measured for ceftriaxone and azithromycin by E-test (BioMérieux, UK) on GC agar base with 1% vitox at 37°C in 5% CO_2_.

### Whole-genome sequencing

Genomes were sequenced by MicrobesNG (https://microbesng.com), with Illumina next generation sequencing using a 250 bp paired end protocol. Some of the 367 isolates (n = 13) were omitted after failure to obtain viable colonies. Sequence read data for the remaining 354 isolates in this study were deposited in the European Nucleotide Archive (ENA) under accession number PRJEB63206. Accessions and quality control statistics are available at github.com/.

### Population structure analysis

Genomes from the G-ToG collection were uploaded to the PubMLST database (https://pubmlst.org/neisseria/) (Jolley *et al*., 2018). Population structure was investigated using 26,802 global *N. gonorrhoeae* isolates publicly available on PubMLST [accessed 11 October 2025] dating from 1940 to 2025, and analysed with the *N. gonorrhoeae* core genome multilocus sequence typing (cgMLST) scheme (v 2), which includes 1,668 core genes (Harrison *et al*., 2020). Minimum spanning trees were constructed in PubMLST using GrapeTree (Zhou *et al*., 2018) with the MSTreeV2 algorithm, based on allelic distances between isolates, and then nodes annotated according to the first two components (superlineage) of the Life Identification Number (LIN) code (Unitt *et al*., 2025).

### Phylogenetic construction and analysis

Sequence reads of 354 *N. gonorrhoeae* clinical isolates from MicrobesNG were mapped to reference strain FA1090 (GenBank accession NC_002946.2) using Snippy v 4.6.0 (Seemann, 2015). The resultant core single nucleotide polymorphism (SNP) alignment was then used to create a recombination corrected maximum-likelihood phylogenetic tree through Gubbins v 3.3.5 (using IQ-TREE v 2.3.5) (Croucher *et al*., 2015, Nguyen *et al*., 2015). Chromosomal resistance determinants (*penA*, *ponA*, *mtrR*, *porB* and 23S rRNA) and gonococcal mobile genetic elements (p*bla*, pConj, pCryp and the gonococcal genetic island (GGI)) were identified using PubMLST. The phylogenetic tree and associated metadata were visualised using Interactive Tree of Life (iTOL) software (Letunic and Bork, 2021). SNP differences between isolates from different anatomical sites within the same participant were calculated using Snippy, where one isolate in each pair was selected as the reference.

### Statistics and statistical modelling

To assess whether gonococcal lineages were structured by membership to a sexual network, the phylogenetic signal of men who have sex with men (MSM) versus heterosexual-associated isolates was quantified using Fritz and Purvis’ D statistic (Fritz and Purvis, 2010) on the core genome phylogeny, with 5,000 permutations using the *caper* package in R (v 4.3.1). D < 0 suggests very strong phylogenetic clustering of isolates with the phenotype.

To examine the factors influencing AMR susceptibility to ceftriaxone and azithromycin, multivariable linear regression models were fitted with log₂-transformed MIC as the outcome variable. Resistance predictor variables included presence of mosaic/semi-mosaic *penA*, *ponA* L421P mutation, *mtrR* promoter deletions, *mtrR* coding sequence mutations, *porB* substitutions at G120+A121 positions and AMR plasmid carriage (p*bla* and pConj carrying *tetM*). Regression analyses were restricted to isolates with complete genotype and MIC data (n = 287/354), and collinearity was assessed using variance inflation factor (< 5 indicating acceptable independence of predictors). Resultant regression coefficients (β) correspond to the estimated change in log_2_ MIC independently associated with the presence of each determinant. Positive β values indicate increased MICs, while negative values indicate a correlation with decreased MICs. All analyses were conducted in R.

Comparisons of MICs between sexual networks were evaluated using Mann–Whitney U tests, while associations between the co-occurrence of mobile genetic elements were assessed using Fisher’s exact tests, with effect sizes reported as odds ratios (ORs) with 95% confidence intervals (CIs). GraphPad Prism (GraphPad Software v 10.6.0) was utilised for these statistical analyses, and significance thresholds were defined as *p* ≤ 0.05, with *p* ≤ 0.001 (***) and *p* ≤ 0.0001 (****) indicated where appropriate.

## RESULTS

### Population structure of N. gonorrhoeae isolates in the G-ToG collection

Genome sequenced clinical *N. gonorrhoeae* isolates (n = 354) from the G-ToG trial (Ross *et al*., 2019), were collected in 2014 (n = 19), 2015 (n = 159) or 2016 (n = 176). To place the G-ToG isolates in global context, a cgMLST minimum spanning tree of 26,802 global PubMLST isolates from 74 countries (Jolley *et al*., 2018) was constructed, revealing extensive diversity with multiple globally disseminated lineages (Fig 1A).

**Figure 1.**
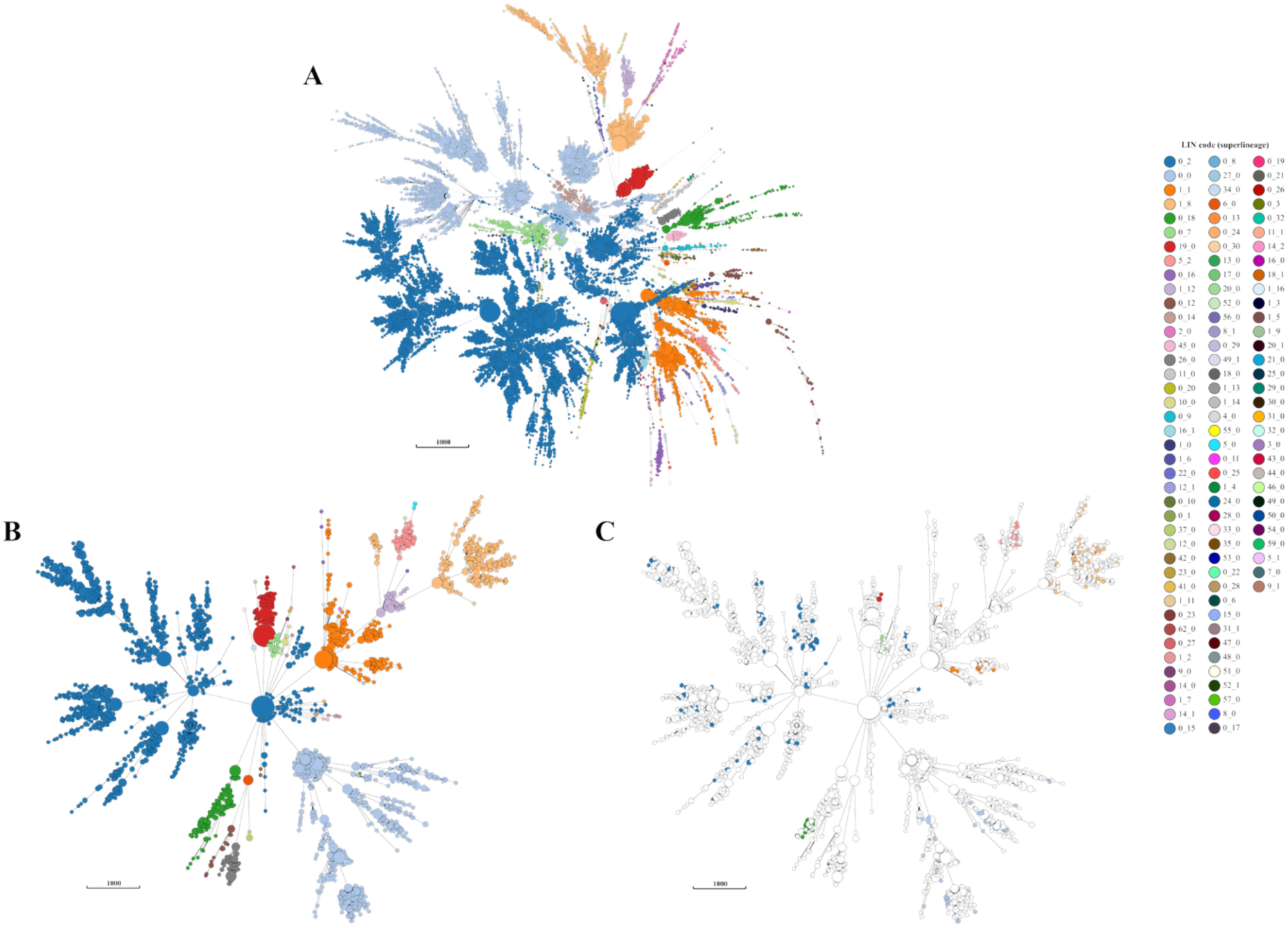
Population structure of *N. gonorrhoeae* isolates. Minimum spanning trees of *N. gonorrhoeae* isolates from the PubMLST (Jolley *et al*., 2018) database were generated based on cgMLST and annotated with LIN code (superlineage). Trees show the population structure of **A)** 26,802 global isolates, **B)** 4,013 UK isolates and **C)** the same selection of UK isolates but 354 G-ToG isolates highlighted only.

A second minimum spanning tree of 4,013 UK isolates revealed most major global lineages circulate in the UK (Fig 1B), indicating the gonococcal diversity in the UK largely mirrors the global population in PubMLST. The structure of the gonococcal population in the UK is consistent with frequent international importation of strains alongside sustained local transmission. When the 354 G-ToG isolates were overlaid on the tree of isolates from the UK and annotated according to their LIN code (superlineage), they were distributed across the tree with no restriction to or over-representation of a particular lineage (Fig 1C). This demonstrates that the G-ToG collection largely captures the breadth of gonococcal genetic diversity circulating in the UK, providing a robust foundation for downstream analyses that is comparable to the wider population.

### Epidemiology of N. gonorrhoeae isolates in the G-ToG collection

The 354 G-ToG isolates were obtained from 296 participants of whom 81.4% (241/296) were male and 18.2% (54/296) were female (Table 1). Amongst males, 74.3% (179/241) were MSM. HIV positivity was recorded in 14.8% individuals, the majority of whom were MSM (95.5%), aligning with other studies showing higher prevalence of HIV among MSM populations (Desai *et al*., 2017). Most participants were white (68.5%), and the modal age group was 25-34 (38.5%). A significant proportion of individuals (41.5%) had been infected with gonorrhoea at least once before and 26.6% of individuals were concurrently infected with chlamydia, diagnosed using a NAAT. Infection site varied by reported sexual behaviour; in heterosexual males, 90.9% (60/66) of isolates were from the urethra, whereas MSM and female participants had infections distributed across genital (51.2%, 147/287), rectal (28.9%, 83/287) and pharyngeal (17.8%, 51/287) sites.

To determine whether participant metadata correlated with gonococcal isolates, a core SNP maximum likelihood tree for all 354 G-ToG isolates was constructed using *N. gonorrhoeae* FA1090 (GenBank accession NC_002946.2) as a reference, then compared to matching clinical and demographic data. No associations were observed between isolate phylogeny and certain participant demographic characteristics, including age, ethnicity, year of isolation or location of the clinic the participant visited (Supp. Fig 1). This suggests that gonococcal lineages within this study are widely distributed across demographic groups with minimal region-specific transmission or significant outbreaks of specific lineages occurred during the study period. HIV positive participants were almost exclusively MSM, and their associated gonococcal isolates dispersed across multiple MSM-associated lineages (Supp. Fig 1). Isolates from participants with chlamydia co-infection also occurred throughout the phylogeny without clear lineage restriction (Supp. Fig 1).

### Multi-site infections within individuals in the G-ToG trial

Of the sequenced isolates within the collection, the majority were obtained from the urethra (46.6%), rectum (23.7%), pharynx (14.7%) and cervix (11.8%). Some participants (16.6%) had multiple gonococcal isolates at different anatomical sites; 40 participants had isolates from two different sites, and nine participants had isolates from three different sites (Table 1). Of those with multiple isolates, almost half of the participants (22/49) had isolates at different sites that were genetically identical (i.e. ≤ 5 SNPs), with 31/49 of participants with isolates ≤ 10 SNPs different (Fig 2B). Interestingly, a small number of the isolate pairs from the same participant (6/49) were genetically highly distinct belonging to different lineages indicated by arrows in Fig 2A, and differing by ≥ 100 SNPs (Fig 2B), consistent with mixed gonococcal infections.

**Figure 2.**
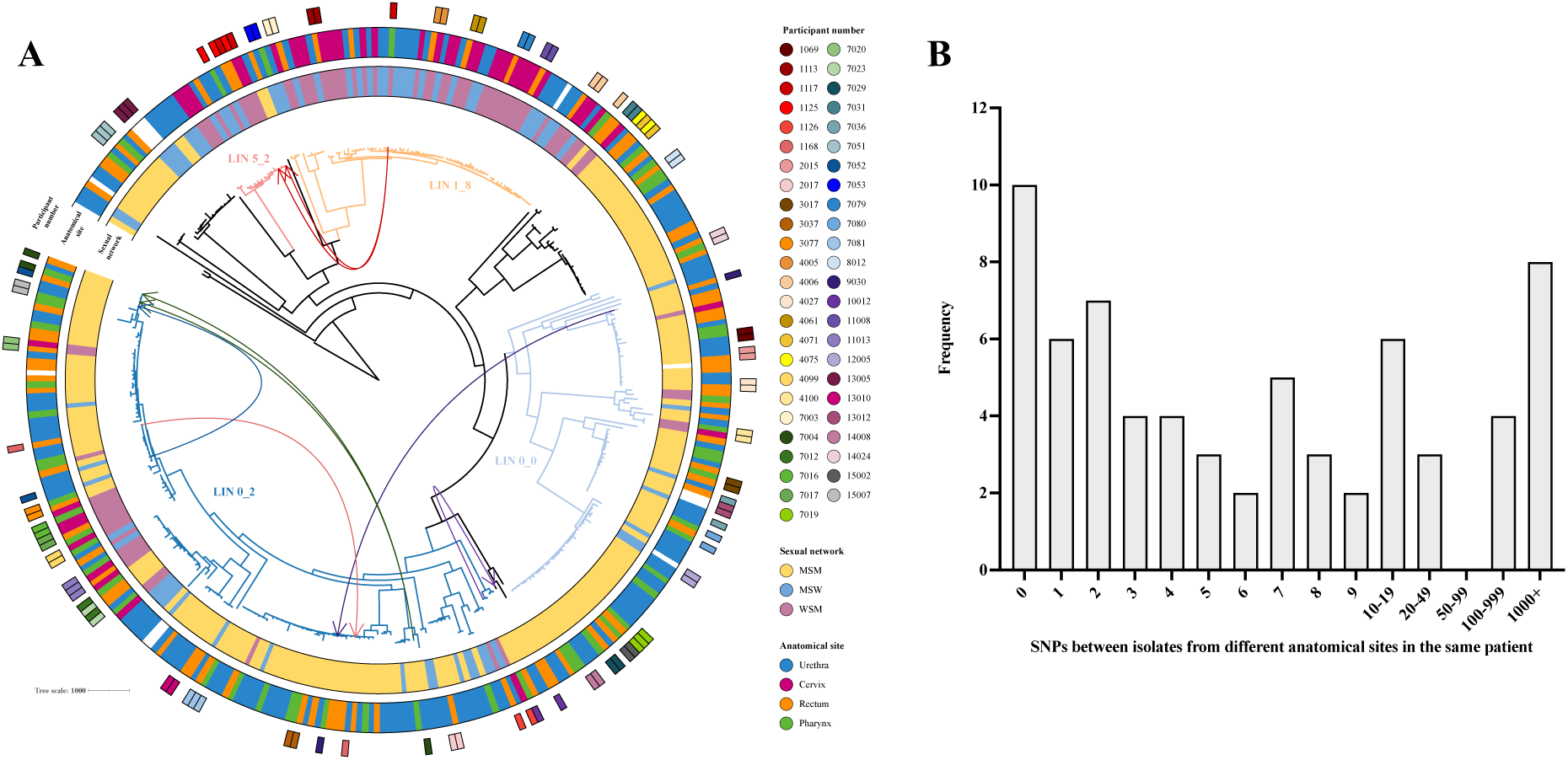
Phylogenetic relationship of *N. gonorrhoeae* isolates from different anatomical sites. **A)** Recombination corrected maximum likelihood phylogenetic tree of 354 *N. gonorrhoeae* isolates from the G-ToG trial mapped to reference strain FA1090, and annotated with the corresponding participant’s sexual network (men who have sex with men (MSM), men who have sex with women (MSW) and women who have sex with men (WSM)) and anatomical site from which the isolate was derived. Participants with gonococcal isolates from multiple anatomical sites are denoted with the participant’s number, with arrows to indicate individuals with genetically distinct isolates. LIN code (superlineage) of several clusters were also highlighted. **B)** Within individuals with multiple-site infections, the frequency of SNPs between each isolate pair was calculated using Snippy (Seemann, 2015).

### Transmission networks shape gonococcal population structure

Numerous epidemiological studies have shown that *N. gonorrhoeae* is predominantly transmitted within MSM networks. In England, 73% of gonorrhoea diagnoses occur in MSM (UKHSA, 2025a) with similar findings in Europe (ECDC, 2025) and the USA (CDC, 2024). Within the G-ToG collection, 60.5% (179/296) of gonorrhoea diagnoses occurred in MSM. When mapped to the phylogeny, membership of a sexual network showed statistically strong phylogenetic clustering across the core genome tree (Fritz and Purvis’ D = 0.19), with the distribution of MSM versus heterosexual (including men who have sex with women (MSW) and women who have sex with men (WSM)) derived isolates differing significantly from random (*p* < 2 x 10^-16^) and showing stronger clustering than expected under a Brownian motion model (*p* = 0.0198). This pattern was driven in part by a large, genetically cohesive clade encompassing LIN 1_8 and 5_2, that was dominated by heterosexual-associated isolates (Fig 3). This clade was comprised of multi locus sequence types (STs) ST1594 and ST11990. This heterosexual clade contained 76.2% (32/42) of all cervical isolates, whereas urethral, rectal and pharyngeal isolates were distributed more evenly across the phylogeny (Fig 2A). Together, these findings indicate that genetically distinct *N. gonorrhoeae* lineages circulate within different sexual networks, with sexual behaviour helping to structure the gonococcal population, despite opportunities for overlap due to male bridging populations.

**Figure 3.**
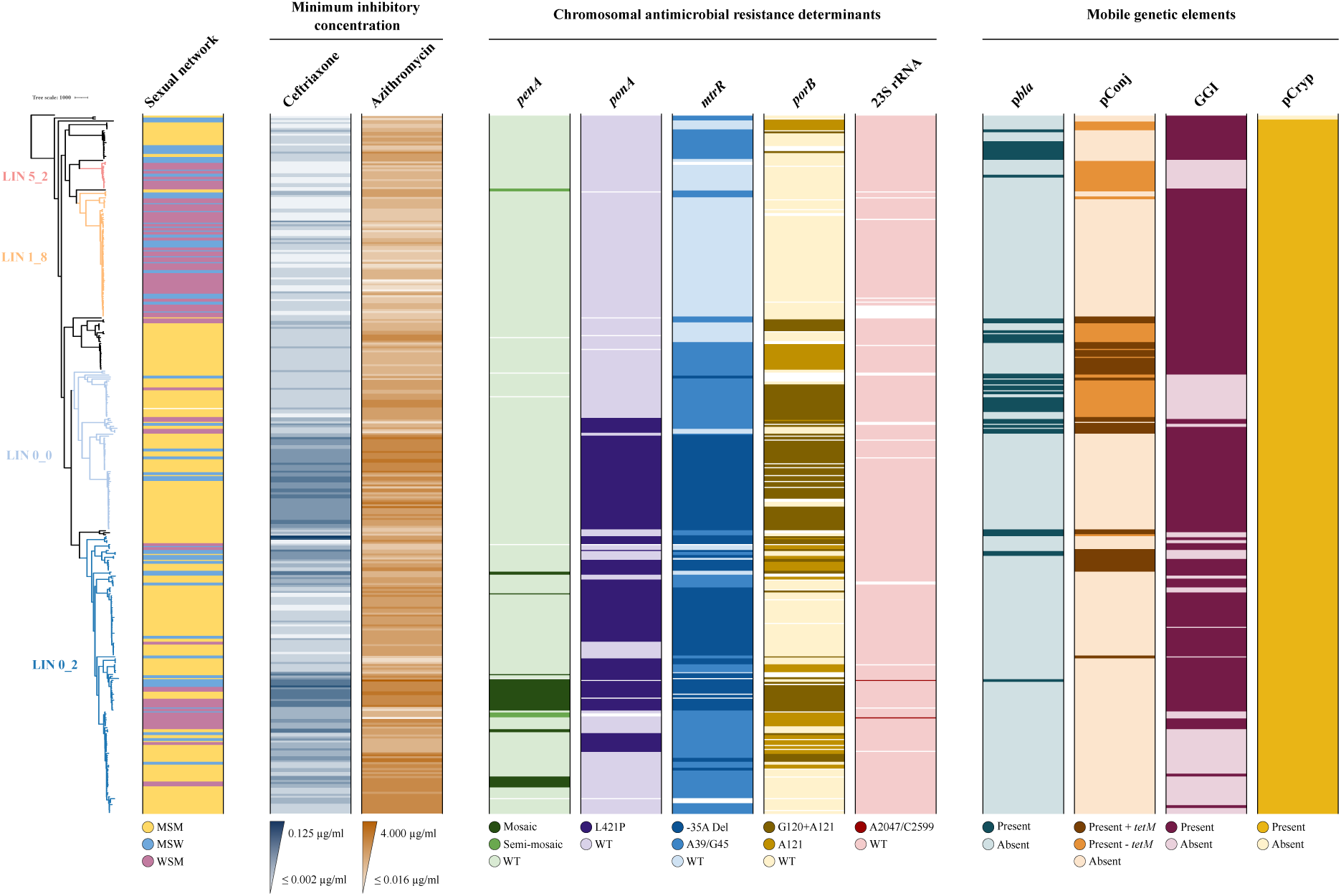
Phylogenetic relationship of *N. gonorrhoeae* isolates with antimicrobial susceptibility and genetic determinants of resistance. Recombination corrected maximum likelihood phylogenetic tree of 354 *N. gonorrhoeae* isolates from the G-ToG trial mapped to reference strain FA1090, and annotated using with the corresponding participant’s sexual network (men who have sex with men (MSM), men who have sex with women (MSW) and women who have sex with men (WSM)), and minimum inhibitory concentration (MIC) from *in vitro* susceptibility testing of isolates to ceftriaxone and azithromycin (Cole, 2020). Chromosomal antimicrobial resistance determinants including *penA*, *ponA*, *mtrR*, *porB* and 23S rRNA were also annotated, with wild-type sequences denoted as WT. Also annotated was the presence of mobile genetic elements, including p*bla*, pConj, pCryp and the gonococcal genetic island (GGI).

### Chromosomal determinants of antimicrobial susceptibility within the G-ToG collection

Key chromosomal loci that underpin the emergence and dissemination of AMR *N. gonorrhoeae* strains include *penA*, *ponA*, *mtrR*, *porB* and 23S rRNA genes. In the G-ToG collection, PBP2 mosaic *penA* alleles were identified in 7.9% (28/354) of isolates (Fig 3), comprising of NG-STAR (Demczuk *et al*., 2017) alleles 34.001, 63.001 and 10.001. Mutations of PBP1 (*ponA*) were common throughout the collection, with 38.4% (136/354) of isolates carrying a L421 substitution (Fig 3). Amongst isolates carrying mosaic *penA* alleles, 64.3% (18/28) also possessed *ponA* L421P. Mutations associated with altered regulation of the *mtrCDE* efflux pump were highly prevalent, occurring in 67.8% (247/354) of isolates (Fig 3). Promoter-region alterations, predominantly the -35A deletion, were identified in 118/354 isolates and coding sequence substitutions in MtrR, principally at positions G45 and A39, were detected in 129/354 isolates. Reduced permeability of PorB and the resulting decrease in β-lactams and azithromycin influx can be mediated by mutations at position A121 and G120 in extramembrane loop 3 (Olesky *et al*., 2002). A121 substitutions (A121S, D, G or N variants) were identified in 144/354 isolates (40.7%), and 94 of these also carried a mutation at G120 (G120K, D or N variants), resulting in 26.6% (94/354) of the collection carrying paired mutations (Fig 3). Mutations in 23S rRNA were rare within the G-ToG collection. Overall, 99.4% of isolates possessed wild-type sequences. Two isolates each carried point mutations previously linked to macrolide resistance (Chisholm *et al*., 2010), A2047G and C2599T (Fig 3).

During the G-ToG trial, the recommended therapy for a gonococcal infection was dual treatment with both ceftriaxone and azithromycin. The susceptibility of all of the isolates to these antibiotics was determined *in vitro* (Cole, 2020). Isolates demonstrated ceftriaxone MIC_50_ and MIC_90_ of 0.004 and 0.016 µg/ml, respectively (range, ≤ 0.002 - 0.125 µg/ml). For azithromycin, the MIC_50_ and MIC_90_ were 0.250 and 0.500 µg/ml, respectively (range, ≤ 0.016 - 4 µg/ml) (Fig 3). To confirm that the observed *in vitro* antimicrobial susceptibility profiles were consistent with established genomic determinants of resistance, multivariable linear regression models were fitted using log₂-transformed MICs for ceftriaxone and azithromycin as outcomes. Both models demonstrated strong explanatory power, accounting for approximately half of the observed MIC variation (adjusted R² = 0.46 for ceftriaxone and 0.47 for azithromycin; full outputs are detailed in Supp. Table 1). For ceftriaxone (Supp Fig 3A), as predicted mosaic *penA* alleles and *porB* substitutions were the strongest independent predictors of increased MIC (β = 0.96, *p* = 1.08 × 10^-5^ and β = 0.99, *p* = 3.3 × 10^-12^, respectively), each corresponding to an approximately two-fold elevation in MIC. In the azithromycin model (Supp Fig 3B), mutations affecting *mtrR* regulation were the dominant drivers of resistance, with promoter deletions (β = 1.94, *p* = 3.4 × 10^-14^) and coding sequence substitutions (β = 1.37, *p* = 4.7 × 10^-12^) associated with substantial increases in MIC, corresponding to a 3.85-fold and 2.59-fold elevation in susceptibility values, respectively. Collectively, these findings demonstrate strong concordance between phenotypic susceptibility testing and known chromosomal resistance determinants for decreased susceptibility to ESCs (Grad *et al*., 2016, Zhao *et al*., 2009) and macrolides (Zarantonelli *et al*., 1999, Hagman and Shafer, 1995).

Azithromycin MICs were significantly higher in isolates from MSM participants compared to isolates obtained from MSW and WSM (Mann Whitney, *p* < 0.0001) (Supp. Fig 2B). Ceftriaxone MICs were significantly higher in isolates from male sexual networks (MSM and MSW) compared to WSM isolates (Mann Whitney, *p* = 0.0002) (Supp. Fig 2A). Clades contributing to this phenotype were those within LIN 0_0, including STs such as ST7363 and ST1570. These associations indicate that isolates within this collection with higher MICs are more frequently observed within MSM-associated transmission networks.

### Distribution of mobile genetic elements in the G-ToG collection

Characterisation of plasmids and their distribution in bacterial populations is key to understanding bacterial evolution and the spread of AMR. For *N. gonorrhoeae*, AMR can be conferred by carriage of p*bla* encoding a TEM-type β-lactamase resulting in high-level penicillin resistance (Elsener *et al*., 2025) and pConj, which frequently carries the *tetM* gene, mediating high-level tetracycline resistance (Cehovin *et al*., 2020).

Within the G-ToG collection, pConj was present in 26.3% (93/354) isolates, with around half of these harbouring *tetM* (39/93) (Fig 3). The p*bla* plasmid was detected in 43/354 isolates (Fig 3). The β-lactamase plasmid requires pConj to mobilise and therefore the plasmids often co-occur (Elsener *et al*., 2025). Co-occurrence of pConj in p*bla* positive isolates was 72.1% (31/43), and 10 of these harboured the *tetM* gene on pConj, which equates to 2.8% of the total collection (10/354) that carried plasmid-mediated resistance to both tetracycline and β-lactams. When compared to the entire collection, p*bla* carriage was significantly enriched among pConj-positive (33.3%, 31/93) compared with pConj-negative (4.6%, 12/261) isolates (Fisher’s exact *p* < 0.0001; OR = 10.4, 95% CI 5.0-21.4). Only 2/354 isolates were found to not carry pCryp, in keeping with published data stating that > 95% of isolates carry this high copy number plasmid (Roberts, 1989), reinforcing its status as a stable plasmid in gonococcal populations.

The gonococcal genetic island (GGI) was widespread throughout 71.2% (252/354) of the collection. There was a significant inverse correlation between GGI and pConj carriage (Fisher’s exact *p* = 2.3x10^-7^; OR = 0.26, 95% CI 0.15-0.42). Given both systems encode type IV secretion machinery, the absence of co-occurrence may reflect selective constraints against maintaining two secretion systems or a mutually exclusive relationship.

## DISCUSSION

WGS of *N. gonorrhoeae* has transformed understanding of gonococcal evolution, population structure and the emergence of AMR (Grad *et al*., 2016, Harris *et al*., 2018, Eyre *et al*., 2017, Golparian *et al*., 2018, Harrison *et al*., 2020). However, most publicly available genomes, including those in PubMLST, are derived from surveillance programmes or outbreak investigations and therefore may not be representative of the broader circulating population. A major strength of the G-ToG trial is the unbiased inclusion of WGS data from all available clinical isolates collected during the trial, coupled with rich and systematically collected participant metadata. This provides a comprehensive and unbiased overview of *N. gonorrhoeae* diversity circulating in England between 2014 and 2016 and enables robust integration of genomic, phenotypic and epidemiological data.

During infection, understanding within-host gonococcal diversity remains an unresolved challenge, as routine clinical laboratories typically retain a single bacterial colony from patients. In this collection, the majority of participants with multi-site infections harboured genetically similar isolates across anatomical sites, consistent with previous reports suggesting limited within-site variation (De Silva *et al*., 2016). Nevertheless, a subset of G-ToG participants carried highly distinct isolates at different anatomical sites, consistent with mixed infections, suggesting that within-host diversity can at times vary considerably between individuals. Incidence of mixed strain infections are likely to be underestimated (Goire *et al*., 2017, Martin and Ison, 2003, Lynn *et al*., 2005) and may have important implications for treatment outcomes, particularly if minor strain populations harbour resistance determinants. Notably, in a separate study using isolates from this collection, genetically near-identical isolates exhibited divergent phenotypes when exposed to the same human serum, with some isolates demonstrating resistance while others remained sensitive (McKeand *et al*., 2026). This indicates that even isolates indistinguishable at the core genome level may differ functionally, potentially due to regulatory mechanisms such as phase variation or differential expression of surface-exposed factors that are not captured by standard genomic analyses. Mixed infections also complicate surveillance and vaccine efficacy assessments, as antigenic heterogeneity across co-infecting strains could influence both individual-level protection and population-level immunity.

Integrating phylogenetics with behavioural metadata has demonstrated that gonococcal population structure of the G-ToG collection is strongly shaped by sexual network membership, with isolates clustering significantly by MSM versus heterosexual transmission groups. This pattern is consistent with current epidemiology data from England (UKHSA, 2025a), which show that gonorrhoea disproportionately circulates within MSM sexual networks.

In the G-ToG study, isolates associated with the MSM sexual network exhibited higher MICs compared with isolates from heterosexual networks. Similar patterns were observed by contemporaneous national surveillance data in the USA during 2014-2016, which found the majority of isolates with elevated azithromycin MICs occurred in MSM compared to heterosexual males (CDC, 2017). In contrast, European surveillance data from the during this period indicate only modest differences in isolate MICs between sexual networks (Town *et al*., 2017, Cole *et al*., 2017, ECDC, 2018).

MSM-associated lineages in the G-ToG collection included globally disseminated multi locus STs such as ST7363 and ST1579 (within LIN 0_0). ST7363 is a successful worldwide clone which has a capacity to develop high-level resistance to ESCs, and ST1579 is a common international lineage that often carries a combination of chromosomal resistance determinants (Lin et al., 2023, Unemo et al., 2012, Harrison et al., 2020). Despite encompassing these STs, all isolates in the G-ToG collection remained within the clinical resistance threshold for ceftriaxone, consistent with reports that clinically relevant ceftriaxone resistance remains rare in England (UKHSA, 2024).

Conversely, a large heterosexual-associated clade (LIN 1_8/5_2) comprised of ST1594 and ST11990, was characterised by minimal resistance determinants and markedly lower MICs to both ceftriaxone and azithromycin. This was supported by Ma *et al*. (2020), where a link was found between cervical isolates and increased antibiotic susceptibility, due to an over-representation of loss of function mutations in the *mtrCDE* operon. ST1594 has previously been documented as typically being susceptible to antibiotics (Ilina *et al*., 2010). A report on isolates from the Netherlands collected during a similar time period stated similar findings, where isolates from ST11990 were only found circulating in heterosexual patients (de Korne-Elenbaas *et al*., 2021), with a significant association with female patients.

This dataset represents *N. gonorrhoeae* clinical isolates collected a decade ago (2014-2016), providing a baseline view of *N. gonorrhoeae* population structure and AMR in England, prior to the introduction of several key public health interventions in the UK. These include the introduction of HIV pre-exposure prophylaxis (HIV PrEP) (UKHSA, 2020) from 2016 and the 2025 roll out of both doxycycline post-exposure prophylaxis (Doxy-PEP) for syphilis control (Saunders *et al*., 2025) and Bexsero vaccination (UKHSA, 2025b) for the prevention of gonorrhoea, which are targeted towards higher-risk individuals. Together, these interventions have the potential to alter the landscape of circulating *N. gonorrhoeae* strains; with potential effects through changes in transmission networks, population level immune-mediated selection and AMR. Repeated antimicrobial exposure through Doxy-PEP may exert selective pressure favouring *tetM*-carrying isolates through the enrichment and persistence of pConj. This may in turn indirectly facilitate the co-mobilisation of β-lactamase-encoding p*bla*. Therefore, further continued genomic surveillance is essential to detect emerging changes in lineage distribution and their structure within sexual networks and AMR trajectories, particularly within MSM populations where the burden of incidence remains highest.

## SUPPLEMENTARY DATA

**Supp. Figure 1.**
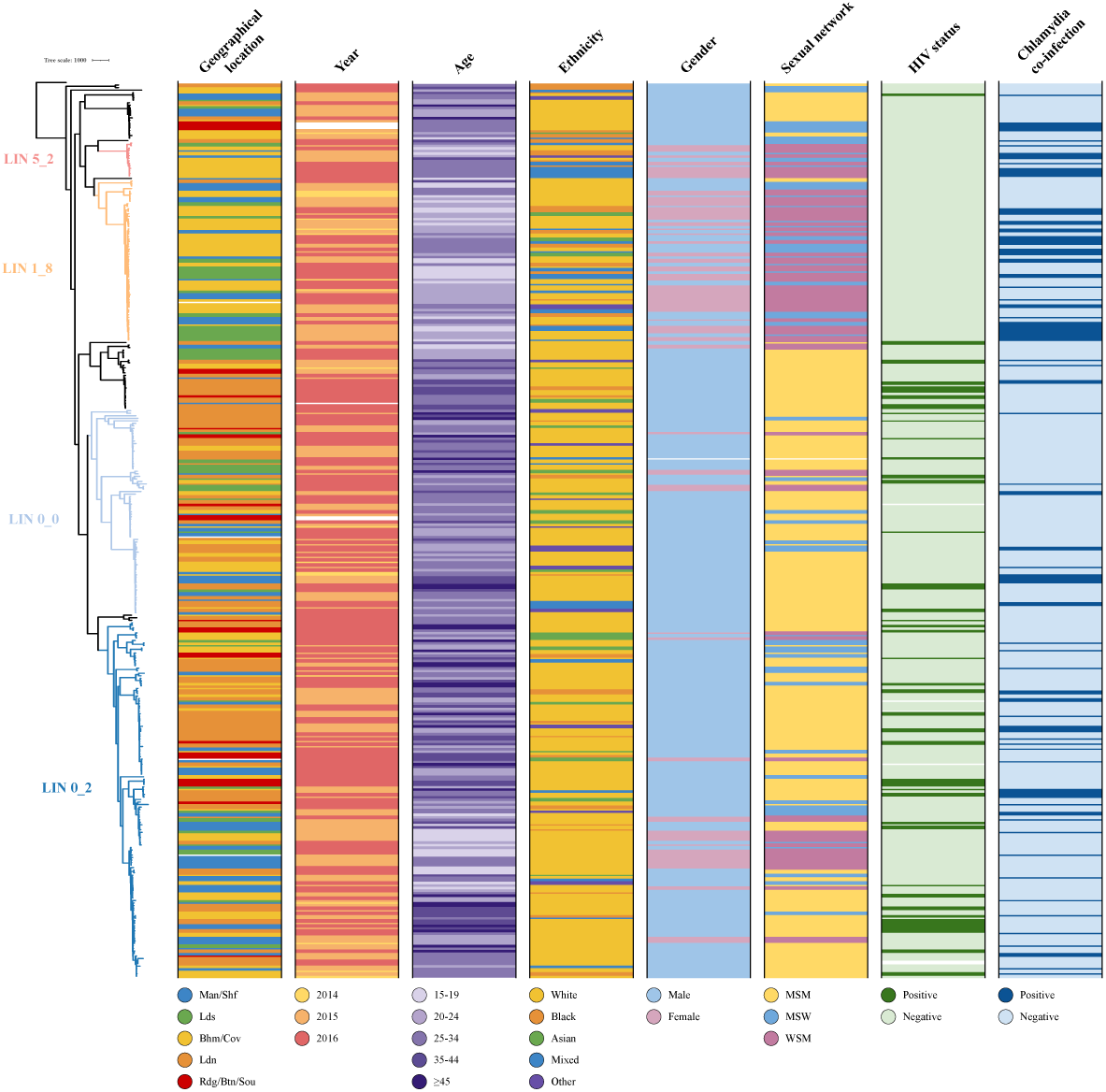
Phylogenetic relationship of *N. gonorrhoeae* isolates with participant demographic data. Recombination corrected maximum likelihood phylogenetic tree of 354 *N. gonorrhoeae* isolates from the G-ToG trial mapped to reference strain FA1090, and annotated with participant demographic data.

**Supp. Figure 2.**
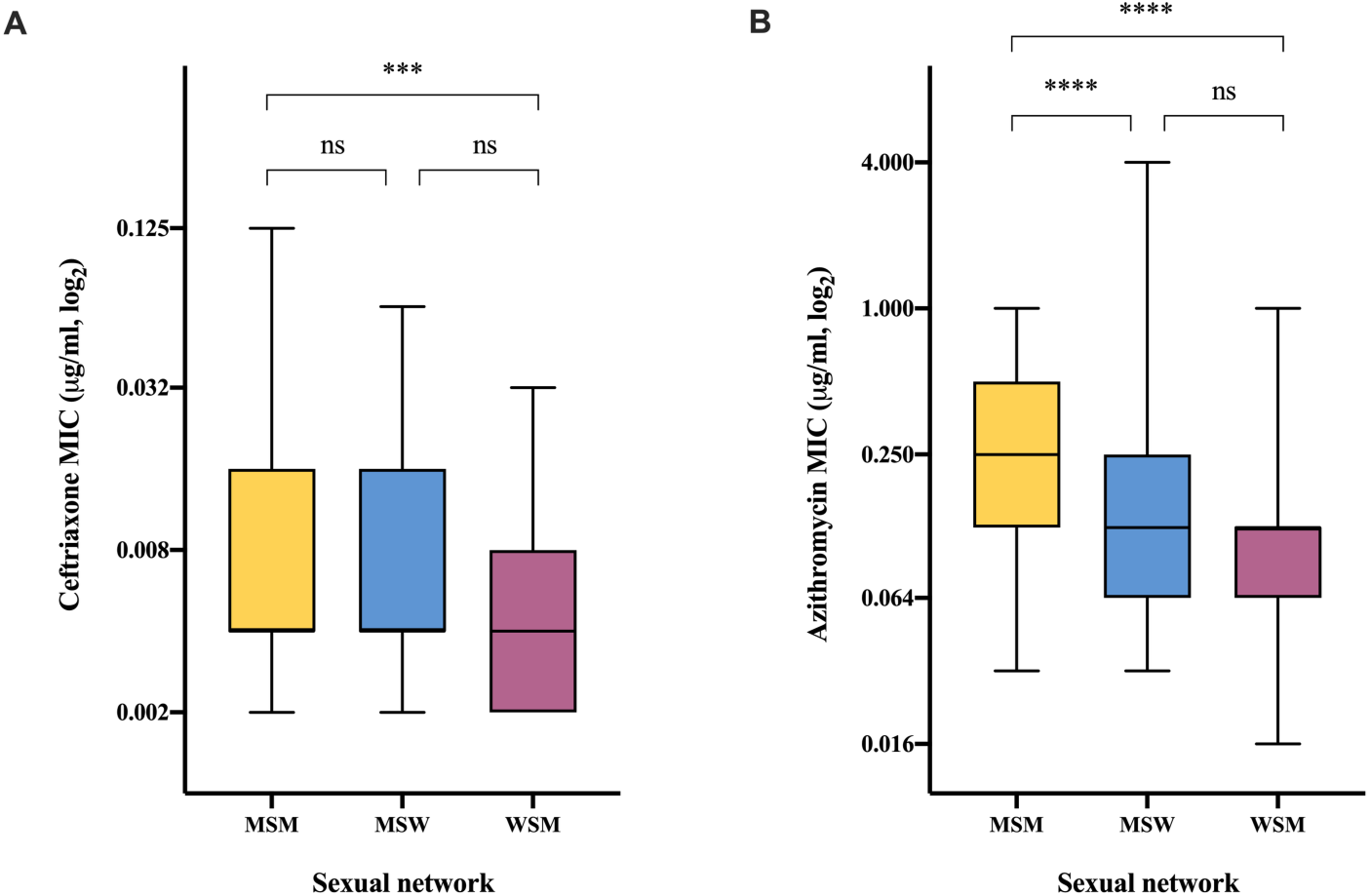
Analysis of MICs of isolates derived from different transmission networks. Comparison of the minimum inhibitory concentrations (MICs) of **A)** ceftriaxone and **B)** azithromycin between different sexual networks; men who have sex with men (MSM), men who have sex with women (MSW) and women who have sex with men (WSM). Box plots show the median (line), interquartile range (box) and minimum/maximum values (whiskers). Pairwise comparisons were measured using a Mann-Whitney test (GraphPad Prism), and statistical significance is indicated as not significant (ns), *p* ≤ 0.001 (***) or *p* ≤ 0.0001 (****).

**Supp. Figure 3.**
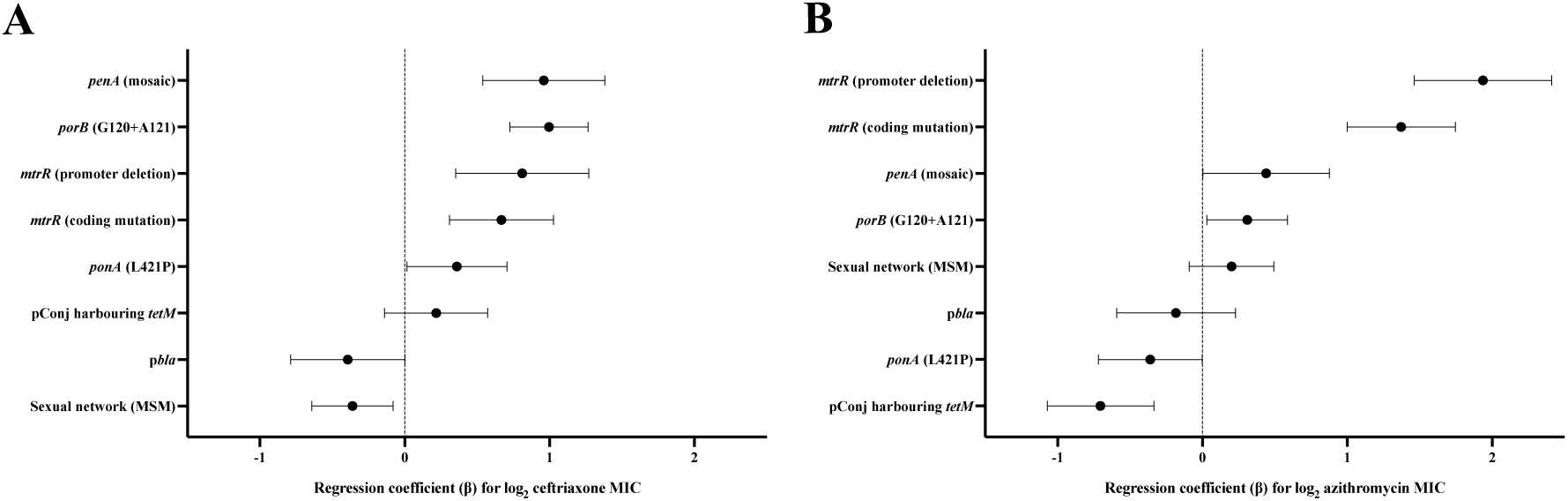
Independent genetic determinants of ceftriaxone and azithromycin susceptibility in the G-ToG isolates. Multivariable linear regression model showing the independent effects of chromosomal and plasmid-associated resistance determinants on log_2_-transformed **A)** ceftriaxone and **B)** azithromycin minimum inhibitory concentrations (MICs). Points represent regression coefficients (β), corresponding to the estimated change in log_2_ MIC associated with the presence of each determinant, with horizontal bars indicating the 95% confidence intervals. Positive β values indicate increased MICs, while negative values indicate decreased MICs. All genetic determinants were included simultaneously in each model. Only isolates with complete genotype and MIC data were analysed (n = 287).

**Supp. Table 1.** Multivariable linear regression models identifying independent genetic determinants of antimicrobial susceptibility in the G-ToG collection. **A)** Ceftriaxone and **B)** azithromycin susceptibility models, fitted using log₂-transformed minimum inhibitory concentrations (MICs) as the outcome variable. Regression coefficients (β) represent the estimated change in log₂ MIC associated with the presence of each genetic determinant, with corresponding standard errors, *p* values and 95% confidence intervals (CIs) shown. An approximate fold-change in MIC is provided for interpretability (calculated as 2^β). Models were restricted to isolates with complete genotype and MIC data (n = 287).

| Determinant | $\beta$ estimate | std. error | *p* value | 95% CI (lower–upper) | Approx. MIC change |
| --- | --- | --- | --- | --- | --- |
| *penA* (mosaic) | 0.96 | 0.214 | $1.08 \times 10^{-5}$ | 0.538 – 1.382 | ~1.94× increase |
| *ponA* (L421P) | 0.36 | 0.176 | 0.0412 | 0.015 – 0.706 | ~1.28× increase |
| *mtrR* (promoter deletion) | 0.811 | 0.234 | $6.05 \times 10^{-4}$ | 0.351 – 1.271 | ~1.75× increase |
| *mtrR* (coding mutation) | 0.668 | 0.183 | $3.17 \times 10^{-4}$ | 0.308 – 1.028 | ~1.59× increase |
| *porB* (G120+A121) | 0.996 | 0.137 | $3.29 \times 10^{-12}$ | 0.727 – 1.264 | ~2.00× increase |
| *pbla* | –0.393 | 0.201 | 0.0513 | –0.788 – 0.002 | ~0.76× (ns) |
| pConj harbouring *tetM* | 0.217 | 0.18 | 0.231 | –0.139 – 0.572 | ~1.16× (ns) |
| Sexual network (MSM) | –0.361 | 0.143 | 0.012 | –0.642 – –0.080 | ~0.78× decrease |

**Supp. Table 1. Multivariable linear regression models identifying independent genetic determinants of antimicrobial susceptibility in the G-ToG collection.**
| Determinant | $\beta$ estimate | std. error | *p* value | 95% CI (lower–upper) | Approx. MIC change |
| --- | --- | --- | --- | --- | --- |
| *penA* (mosaic) | 0.44 | 0.222 | 0.0482 | 0.003 – 0.877 | ~1.36× increase |
| *ponA* (L421P) | –0.361 | 0.182 | 0.0484 | –0.719 – –0.002 | ~0.78× decrease |
| *mtrR* (promoter deletion) | 1.937 | 0.242 | $3.39 \times 10^{-14}$ | 1.464 – 2.410 | ~3.85× increase |
| *mtrR* (coding mutation) | 1.372 | 0.19 | $4.73 \times 10^{-12}$ | 0.999 – 1.745 | ~2.59× increase |
| *porB* (G120+A121) | 0.309 | 0.142 | 0.0301 | 0.030 – 0.587 | ~1.24× increase |
| *pbla* | –0.183 | 0.208 | 0.38 | –0.592 – 0.227 | ~0.88× (ns) |
| pConj harbouring *tetM* | –0.705 | 0.187 | $2.02 \times 10^{-4}$ | –1.071 – –0.336 | ~0.61× decrease |
| Sexual network (MSM) | 0.201 | 0.148 | 0.176 | –0.091 – 0.492 | ~1.15× (ns) |

## Notes

### Competing Interest Statement

The authors have declared no competing interest.

